# Development of Deep-Learning Models that Predict Quantitative Protein-Ligand Interactions in Glycobiology as a part of a Capstone Course

**DOI:** 10.64898/2026.06.19.733466

**Authors:** Haoyan Yin, Weibo Liu, Weihang Zhou, Zechen Chang, Eric J. Carpenter, Aashish Satyajith, Simatsidk Haregu, Russell Greiner, Ratmir Derda

**Affiliations:** Department of Computing Science, University of Alberta, Edmonton, Alberta, T6G 2E8, Canada; Department of Chemistry, University of Alberta, Edmonton, AB T6G 2G2, Canada; Alberta Machine Intelligence Institute (AMII), 1101-10065 Jasper Avenue, Edmonton, Alberta, T5J 3B1, Canada

**Author notes:** These authors contributed equally (Haoyan Yin, Weibo Liu, Weihang Zhou, Zechen Chang).

## Abstract

Glycans coat the surface of all cells, and every glycan is recognised by specific glycan-binding proteins (GBPs). There are no general tools that can accurately estimate the binding strength between glycan and GBP from the amino acid sequence of the GBP and the molecular structure of the glycan, represented as SMILES string. We describe models for predicting such binding strengths developed as a part of a Capstone Course at the University of Alberta. The models are trained on a dataset that combines BindingDB, a published database of small-molecule protein interactions, and data from glycan arrays measured by Consortium of Functional Glycomics (CFG). In this hybrid dataset of protein-ligand interactions the ligands are both glycans from CFG and small molecules from BindingDB; similarly, proteins include GBP and proteins from BindingDB. Three models are presented (i) ProMax which fuses ESM-2, MolFormer, and MolCLR features; (ii) APEX which constrains learning to a predetermined form, a physical model of binding; (iii) UltraMax adds inter-atomic distances for the ligands. To address the dataset’s severe long-tail distribution, the models employ tail-aware losses for rare high-binding instances. Trained and evaluated on approximately one million protein–ligand pairs using hold-out splits for unseen molecules, the three models provide a unified framework for quantitative glycan–protein binding prediction. We observed that learning glycan-protein binding is harder than the similar task of learning small-molecule-protein interactions. Simple mirror-inversion tests led us to postulate that insufficient use of chiral features is an important source of difficulty in learning these interactions.

## Introduction

This work began as a project in the Computing Science “Artificial Intelligence Capstone” course CMPUT 469 at the University of Alberta. Two domain experts, working in glycomics, along with two machine learning (ML) researchers—a professor and a graduate student teaching assistant —guided a team of four senior undergraduate students on a four month long real-world research project: to build an ML model that can predict whether a protein will bind to a glycan or other molecule. The Capstone project involved first identifying the task itself, then iteratively addressing relevant issues with feedback from a domain expert, leading to an implementation and culminating in evaluating that system.

Glycan–protein interactions are important in cell–cell communication, immune recognition, host–pathogen attachment, and disease progression ^1,2^. Predicting these interactions is challenging because glycans are structurally diverse branched oligomers of monosaccharides whose binding depends on composition, position, and stereochemistry ^2,3^. Prior work – including BioLK, CCARL, GlyNet, CarboGrove, and LectinOracle ^4-8^ – show that ML can effectively learn models that can predict the binding strength between glycans and GBPs. Of course, GBPs are merely a subset of proteins; similarly, glycans are merely one subset of molecular ligands. Separating glycan-GBP interaction from the general task of predicting protein-ligand interactions led to challenges, such as the scarcity of data describing binding of GBP to glycans and even uncertainty about what constitutes a *bona fide* glycan or GBP. Foregoing historical separation, we combine glycan-GBP data with data for other protein-ligand datasets, where the term “ligands” designates any molecular interacting partner of a protein that has a moderate size can be accurately represented using a universal all-atomic representation, such as SMILES ^9^.

To enable machine learning from the expanded protein-ligand dataset, the estimation of protein-ligand binding strength should be reduced to a universal binding parameter. For example, Bojar et al. ^5^ have shown that Relative Fluorescence intensity Units (RFU) measured on glycan arrays can be reduced to parameters like Z-scores (Z′) to train ML models. Such Z′ are used universally in high-throughput screening of binding interactions of all classes of molecules. Our previous publication converted the same RFU to a fraction bound parameter, *f*, to blend RFU with other measurement endpoints ^10^. For example, *f* is trivially computed from any binding parameters like equilibrium dissociation constants (K_D_) or half-inhibitory concentrations (IC_50_) ^10^. We demonstrated that glycan data converted to SMILES and *f* derived from RFU can be used for training of the MCNet model with performance analogous to all previously reported models ^10^. Although Z and *f* are similar algebraic forms of binding observations, the explicit calculation of *f* is more productive because such data is more intuitively related to *bona fide* fraction bound values derived from thermodynamic values such as *K*_D_: in case of simple equilibrium P + L → PL, between protein P and ligand L, the *f* = [L] / ([L] + [PL]), where square brackets denote equilibrium concentrations. In 1:1 binding equilibrium, the Henderson-Hasselbach equation explicitly demonstrates that only [P] is necessary and sufficient to compute the *f*^10^ (Fig. S2). The same formalism can be applied to the immobile ligand on the surface of glycan array, and again, concentration of soluble protein [P] is necessary and sufficient to define *f* = [L*] / ([L*] + [PL*]), where stars denote the immobile ligand tethered to the solid surface or its complex with protein

Representation of glycans is a central challenge in computational glycobiology. Unlike proteins and nucleic acids, which have linear structures, glycans are built-up of monosaccharides in a branched, tree-like structure. Many natural glycans are related as stereoisomers and learning to distinguish properties of molecules with identical compositions but different chiralities for specific atoms is reported to be one of the hardest problems for contemporary ML models ^11^. Experimental analysis of natural glycans often reports structures with uncertainty in chemical or stereochemical compositions. Databases such as GlyTouCan and GlyGen embrace such uncertainties and employ specialized glycan notations such as WURCS ^3^ to represent glycans with such uncertainties. For instance, of the 253,779 entries in GlyTouCan ^12^ only 39,030 are annotated as fully-defined. Glycans whose stereochemical and compositional aspects are all defined, can be represented in universal ML-compatible formats such as canonical SMILES strings, using tools such as GlyLES ^13^. Other ML-compatible descriptions of glycans use hierarchical approaches that combine mixed atom-level and monosaccharide-level embeddings ^14-16^. We do not pursue these approaches because they make it difficult, if not impossible, to combine glycan data with general small-molecule data that has obvious representation as monomers/oligomers.

Early computational work on protein–glycan binding emphasized motif discovery from glycan microarray data: examples are GlyGen-style analysis pipelines and subtree-mining approaches for identifying recurrent glycan determinants ^6,17^. CarboGrove extended a traditional motif discovery/analysis pipeline by fitting binding curves to interactions while also incorporating information such as protein concentration, and glycan display densities ^8^. GlyNet established a quantitative multi-output regression benchmark for protein–glycan interaction prediction on glycan-array data ^4^, MCNet models merged glycan array RFU values with *K*_D_ values of glycomimetic molecules ^10^, and LectinOracle combined protein transformers with glycan graph encoders to improve generalization across new lectins and glycans ^4,5,7,18^. This progression from motif-centric analysis to learned multimodal models motivated our own design. We drew inspiration from CarboGrove ^8^, SweetNet ^19^, GlyNet ^4^, MCNet ^10^ and LectinOracle ^5^ and noted room for improvement: GlyNet and MCNet are primarily glycan-driven against a (predetermined) panel of protein samples. LectinOracle does not explicitly target concentration-conditioned regression and cannot input general small molecules that are not oligomers of monosaccharides. More generally, prior work often emphasizes specialized glycan encoding. Our research tested the plausibility of an expanded multimodal architecture that jointly encodes protein sequence, concentration, and glycan structure from universal SMILES strings, modeling their interactions through gated fusion and cross-attention for continuous prediction of binding. We adopted pretrained encoders and learned fusion layers to model interaction strength directly rather than relying only on hand-crafted descriptors or motif rules.

This manuscript describes an end-to-end deep learning pipeline, consisting of three stages: data preprocessing, multimodal feature extraction using biochemically informed encoders, and interaction modeling with progressively more expressive architectures. Many of the molecule-protein pairs have near-zero fraction-bounds (i.e., do not bind) and to manage the label imbalance we also introduce tail-aware objective functions to handle the long-tailed distribution of binding data. The methodology section describes the datasets and the model architectures. We then evaluate the performance of the models and show sample predictions across protein, molecule, and concentration input axes. We show that models that are not trained on stereoisomers insufficiently utilize chirality information even though the molecular encodings present this information to the model. This leads to similar predictions between a chiral molecule and its mirror image (enantiomer), even though if there is binding in the former case, it is expected to be ablated.

## Results

### Analysis of the dataset

The dataset consists of approximately one million protein–glycan interaction pairs from CFG ^20^ and protein-molecule pairs from BindingDB ^21,22^. The interaction pairs from BindingDB contain glycan-like molecules as well as other small-molecules (non-glycans). Including these in the training data is expected to enhance the ability of the models to learn and predict the behaviours of a broader chemical family, enabling the model to handle more chemically diverse glycans than the ones on which it is trained, as well as helping to support learning on the spacer/linker molecules used to attach the glycan in question to the assay in many glycan-protein experiments including the CFG data.

Not all the data in the datasets was appropriate for our use. For BindingDB, the available *K*_D_ values were filtered to only include proteins for which a minimum of 30 molecules with *K*_D_ values exist, or alternatively a minimum of 30 proteins for a molecule. These values were then converted into fraction bounds at concentrations of 0, 0.1, 1.0, 10, 100, …, 10^5^ nM. Note that 44,794 K_*D*_ values are not specific numbers, but are instead limits, e.g. >10000 nM or <0.2 nM. These values still convey significant information about the corresponding molecule-protein interactions and we intentionally sought to retain these in the data. Accordingly, we take these to have fraction bounds of zero below the minimums and one above the maximums, but because we do not know them, we do not produce fraction-bounds outside of these intervals. The 110,172 *K*_D_ values that we use from BindingDB represent 75,561 unique molecule-protein pairs, of which 54,534 pairs only occur once. Some 1,098 molecule-protein pairs across 9,791 *K*_D_ values are highly replicated - each having five or more *K*_D_ values. Similarly, not all CFG ^20^ data was included, as we only used instances that reported protein concentration and the protein sequence. The CFG data also has duplicate values, of the 758 array experiments used, 495 (65%) were from unique protein-concentration pairs, with ten highly repeated protein-concentration pairs (68 arrays) occurring five or more times.

We did not attempt to reconcile the replicate values in either source dataset, either by averaging them or by suppressing data points, instead passing all of them into our fraction bound calculations, and subsequently into the models, which then must learn in the presence of the noisy and inconsistent training data. Many of these replicate values will show minor variations associated with experimental noise, but some of these numbers have large differences, sometimes of several orders of magnitude; see Fig. 1F for distributions of the standard deviations. We consider that a reasonable level of uncertainty in these numbers is a standard deviation of less than one order-of-magnitude. After introducing a threshold at this level, we observe that only 2716 cases (13%) of the data points violate it among the 21,026 multiple datapoints. Since the remaining 87% of the multiple datapoints are consistent with each other, any machine learning model capable of accurately predicting fraction bound can be expected to interpolate between these values but will nevertheless have a baseline error (explored later in Fig. 3).

**Fig. 1.**
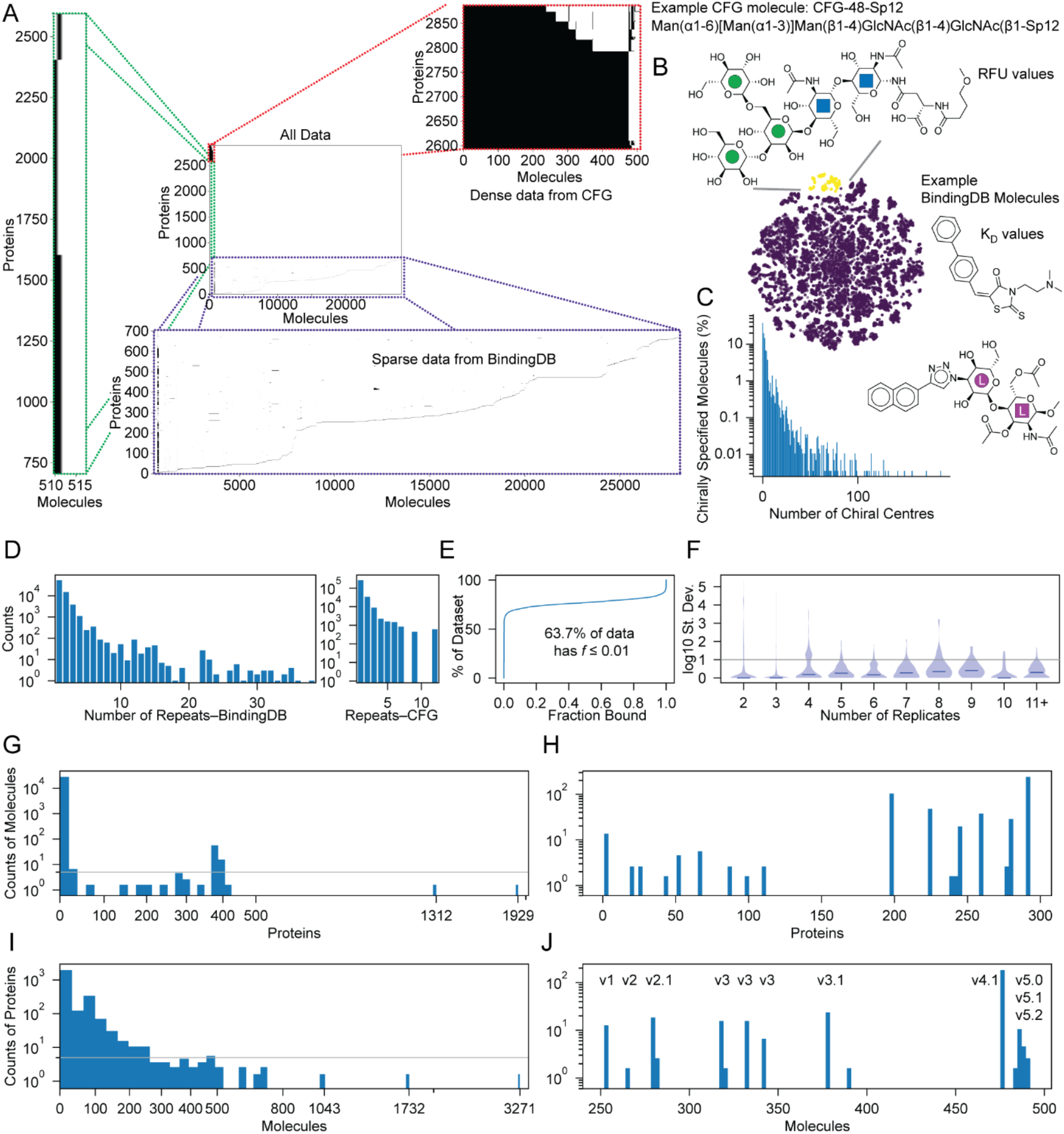
Data sources and analysis. A) Data is drawn from CFG and BindingDB. While the CFG dataset is dense with data for most of the glycan-protein pairs (red box), the BindingDB data is sparse with only a few of the possible molecule-protein pairs (blue box). B) The molecular structures include spacer/linkers for the CFG data and include glyco-mimetic molecules from BindingDB. A t-SNE plot of the molecules shows that the glycans in the CFG set occupy only a small fraction of the total molecular space (yellow), with the BindingDB molecules (purple) filling nearly the whole space. C) An important property of glycans is the large number of chiral centres in each monosaccharide. We note that many of the molecules in our set have many chiral centres. D) The number of experiments corresponding to different protein-molecule pairs in BindingDB and CFG. E) About 2/3rds of the instances are non-binding, creating a so-called long-tailed distribution. F) When the same molecule-protein pair is repeated in BindingDB, the reported K_*D*_s are often very close, even identical, but some have orders-of-magnitude differences. Distributions of the replicate dispersions vary with the number of replicates. The medians of the distributions are shown (horizontal line). G) Most proteins taken from BindingDB are only paired with a small number of molecules, in many cases under 5 (horizontal line), while those from CFG H) have distinct patterns. I) Similarly, molecules from BindingDB are only present in a small number of data cases, while J) those in CFG show distinct clumps associated with particular array versions.

In line with standard ML practice, we split the molecules into train/test/validate subsets, herein using an 70%/20%/10% division. The test subset is completely excluded from the training optimization and then the models are evaluated on it. Included in the test set is a group of 136 molecules drawn from CFG, see Fig. 1. The molecules in this group are primarily glycans composed of monosaccharides as well as lactose variations, including with and without sialylation, and human ABH-blood group antigens.

### Protein and molecule encodings

We adopt pretrained models that provide latent representations for both protein and molecules. Protein features were obtained from ESM-2 embeddings ^23,24^, while glycans and BindingDB molecules were represented using MolFormer embeddings ^25^, MolCLR molecular topology-aware embeddings ^26^, and ETKDG-modelled inter-atomic distances ^27^. ESM-2 has an intrinsic maximum length of 1022 amino acids. As an initial approach, this was handled in the simplest possible way, by truncating longer protein sequences to only the first 1022 amino acids before embedding. This affects 11% (363) of the protein sequences. The ESM family produces an embedding describing each amino acid in the context of the other residues in the sequence. The embeddings corresponding to each residue in the protein is updated based on the whole sequence as they are passed through attention blocks, and hence each residue’s embedding contains information about the whole protein. The mean of these is commonly used to produce a final representation of the whole protein; instead, we retained the embeddings of the sequence from the first 256 residues (shorter sequences were zero-padded). This produced a more detailed representation of each protein (relative to the averaging approach) in a fixed-size array of 256 x 1,280 features using the esm2_t33_650M_UR50D model.

Molecular encodings constructed from multiple modalities were used to represent glycans and small-molecules. MolFormer is presented with SMILES strings and relies on its transformer to reconstruct relationships between different parts of the string; such an approach may capture global molecular contexts ^25^. MolCLR converts SMILES strings into atom-level graphs and its encoder operates from these molecular topologies ^26^. RDKit is used to calculate various physical properties of atoms such the element, hybridization state, number of bonds, charge, and whether it is in a ring, as well as generating a conformer of the molecule from which physical inter-atomic distances are extracted ^28,29^.

### Model descriptions

ProMax (Fig. 2A) is our core multimodal baseline for glycan–protein interaction prediction. The model integrates protein sequence, molecular structure, and concentration through modality-specific encoders, followed by explicit interaction modeling and protein concentration-dependent prediction. Protein sequences are encoded using pretrained ESM-2 embeddings, while molecules are represented through two complementary views: MolFormer embeddings that are hoped may contain global molecular contexts and MolCLR embeddings that are used to encode local graph-based molecular features such as atom-bond connectivity, and substructure-level patterns within each compound. These molecular representations are projected into a shared latent space and fused using a learned gating mechanism. Although ProMax incorporates concentration as an input when directly predicting fraction bound, the model does not explicitly constrain the output to follow the expected concentration-dependent saturation behavior of molecular binding. In contrast, the APEX variants model fractions bound as sigmoidal functions of concentration: instead of directly regressing the fraction bound, *f*, APEX predicts log(K_*D*_) or a latent log(EC_50_). These values are then transformed into the predicted fraction bound at various concentrations using the Henderson-Hasselbach equation. Two variants of APEX were developed. APEX Base predicts log(K_*D*_) values using gated MolFormer embeddings, one-way ligand-to-protein cross-attention, and another gated interaction module. APEX Improved replaces the cross-attention component with self-attention over concatenated ligand/protein tokens. UltraMax (Fig. 2B) extends multimodal binding models by retaining the same ESM-2 protein representations and Mol-Former molecule encoder used in prior versions, while replacing the MolCLR encoder with alternative graph-based representations. It has four main contributions: an explicit bond-graph representation intended to supplement the implicit representation internal to MolFormer; a complementary inter-atomic distance-aware structural branch; an extended gating mechanism that learns jointly over the MolFormer, explicit graph, and distance model; and a deeper bidirectional transformer cross-attention module for protein–molecule fusion.

**Fig. 2.**
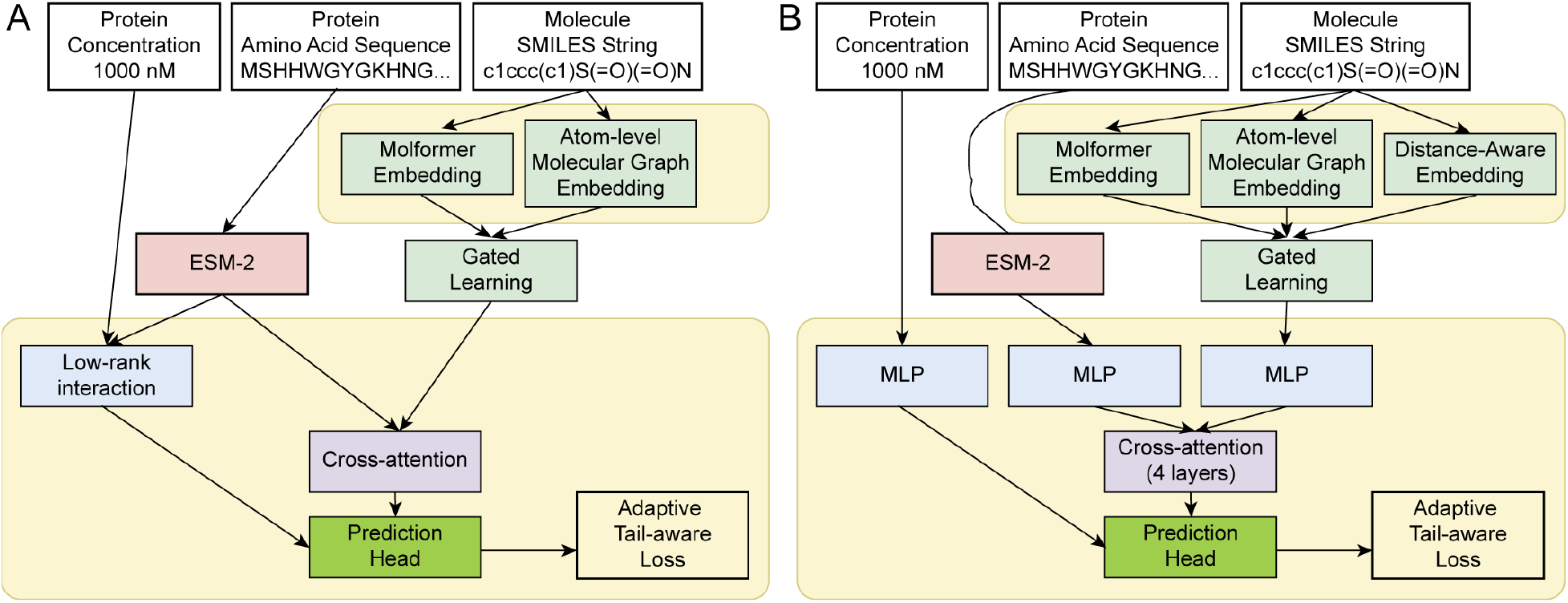
Overview of the ProMax and UltraMax architecture. (A) Protein, molecule, and concentration inputs are encoded separately. Molecule representations from MolFormer and MolCLR are fused through a learned gating module, and the resulting multimodal features are combined through cross-attention and interaction modules before being passed to the prediction head. The model is trained using an adaptive tail-aware loss. (B) In Ultra, the molecule is encoded through three complementary branches: MolFormer embedding, explicit atom-level graph embedding, and distance-aware embedding. A gated learning module adaptively combines these molecule representations. The fused molecule representation is integrated with the ESM-2 protein features and the concentration context through a cross-attention module, followed by a prediction head for final binding estimation.

**Fig. 3.**
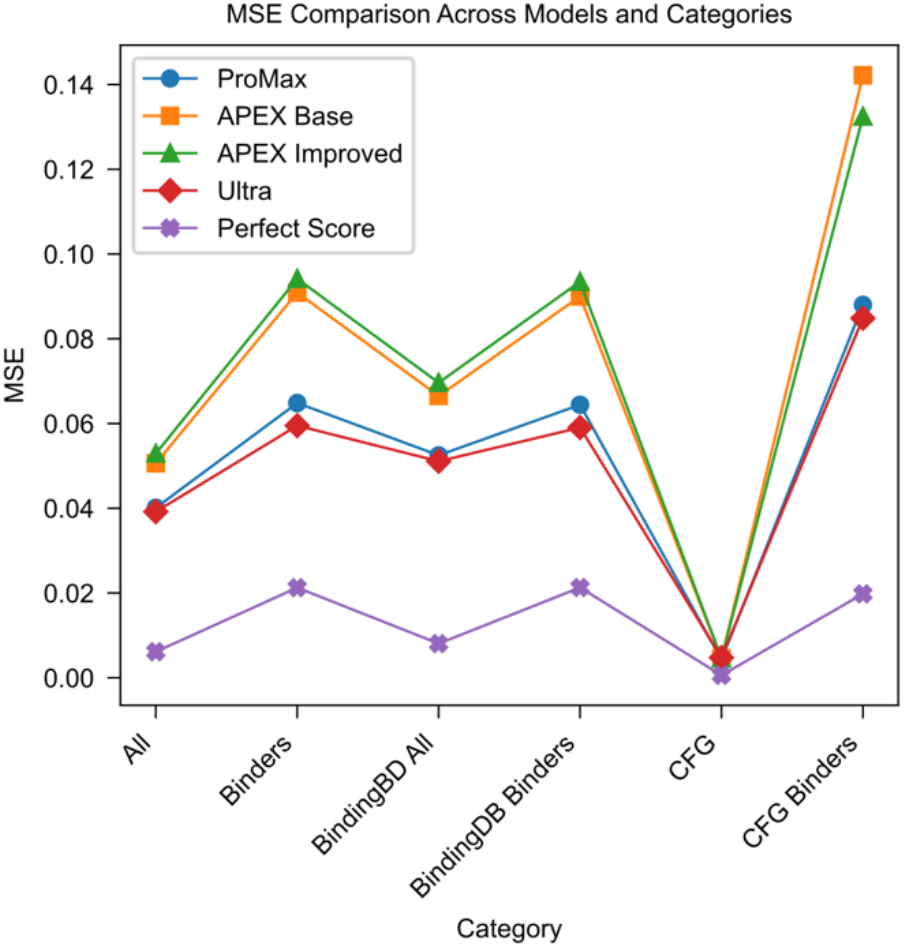
Performance metrics of the four models. Test mean squared error (MSE) across the four models considered. We stratified the MSE based on whether the test instances were from BindingDB or CFG, and whether they were strong binders (fraction bound > 0.2) or not. The data itself imposes some intrinsic variance arising out of inconsistencies in replicate fraction bound entries from the same molecule, protein, and concentration measurements; this variance is reported as the “perfect score” and is a lower bound for any model’s mean squared error.

Examination of the binding data shows that many of the molecule-protein pairs have low fraction-bounds (do not bind) especially at lower protein concentrations (Fig. 1E). To train the ML models in the presence of this long-tail distribution of binding data, we adopt a unified loss that incorporates several ingredients beyond the basic mean squared error (MSE) loss. Most important of these is a second loss term that includes additional penalties for errors where the ground truth is a strong-binder but the prediction is not, and a ranking term that helps the models preserve the relative order or pairs within the model outputs. The APEX models differ from the others in that the ML steps operate in the log(EC_50_) space. Accordingly, in addition to the above losses that are applied after the model output has been transformed to fraction bound values, the APEX models use an additional loss term that operates directly on the log(EC_50_) values. The models were implemented using the PyTorch library. The Supplementary Material provides more detailed descriptions of the models.

### Evaluating model performances

We evaluated the mean squared error (MSE) on the fraction bound predictions across the test set. Given the substantial differences in the molecules from BindingDB and CFG, we confined ourselves to study only the unseen molecule scenario, and stratified our analyses of MSE. We divided the molecules based on their source, the BindingDB dataset versus the CFG dataset. A further division was between molecules with strongly-binding test instances (cases where fraction bound ≥ 0.2 at any concentration). This second stratification is useful since around 80% of the instances in the combined datasets have low fraction bounds, and so a trivial model that always predicts no binding will achieve a low MSE but is not useful. The subsets of the data arising from these divisions were used to evaluate the models. Besides the models, we also considered the theoretically optimal performance in which the mean of the training data for each input (protein, molecule, concentration triples) is always returned as a reference for comparison.

Fig. 3 shows plots of the MSE achieved by the four models in the various categories. Across all data sources (BindingDB, CFG, and both taken together), the models gave worse MSE on the strong binder subset than when all the test data points from that source are considered. The errors across strong binders from CFG were 40% to 65% higher than the errors obtained for strong binders from BindingDB. This observation suggests that it was more challenging for the models to learn glycan-protein binding than more general small molecule-protein binding. In contrast, all models achieved a low MSE on weak-binding CFG examples, demonstrating that this is not a problem with the models learning over the CFG molecules in general.

Comparing across the models, UltraMax had the best performance across all categories, followed by ProMax with physical binding model-constrained APEX models being consistently the worst. Apex Base and Apex Improved had around 25% and 35% higher MSE respectively than the UltraMax model. This is partly because APEX interpolates between one set of data points when multiple sets do not agree on the fraction bound values. We note that APEX Improved gives slightly better performance than APEX Base when it comes to CFG binders, but performs worse in all other categories.

### Sample predictions across molecule, protein and concentration axes

We assessed how the quality of the model predictions compares to the existing variance in the data across molecule, protein, and concentration configurations (Fig. 4A). For ProMax and UltraMax models, the per-configuration MSEs fall close to the per-configuration variance. To understand the behaviour of these plots better, we used the models to predict binding while sweeping through a range of concentrations, giving rise to a curve. Naïvely, we would expect that curve to follow a sigmoidal form, with the onset of binding as the concentration rises above a characteristic threshold. We show some examples of such concentration sweeps in Fig. 4B-C. Notably, in the C case, we observe that the input data (markers) appears to be sampled from two different curves. We see how the ProMax and UltraMax models attempt to deal with this incon-sistency in the data: these models follow the stronger binding data at lower concentrations before shifting to the second set of data points at higher concentrations. The APEX models can only output a single sigmoidal curve and should prefer to output predictions closer to curves supported by more points. This may explain why the APEX model prefers the leftmost curve in Fig. 4C, middle two panels, as there are more data points consistent with this solution than with the rightmost curve. An interesting case occurs when there are two EC_50_ values that lead the APEX models to have an MSE close to twice the lower-bound value (Fig. 4A, middle two plots).

**Fig. 4.**
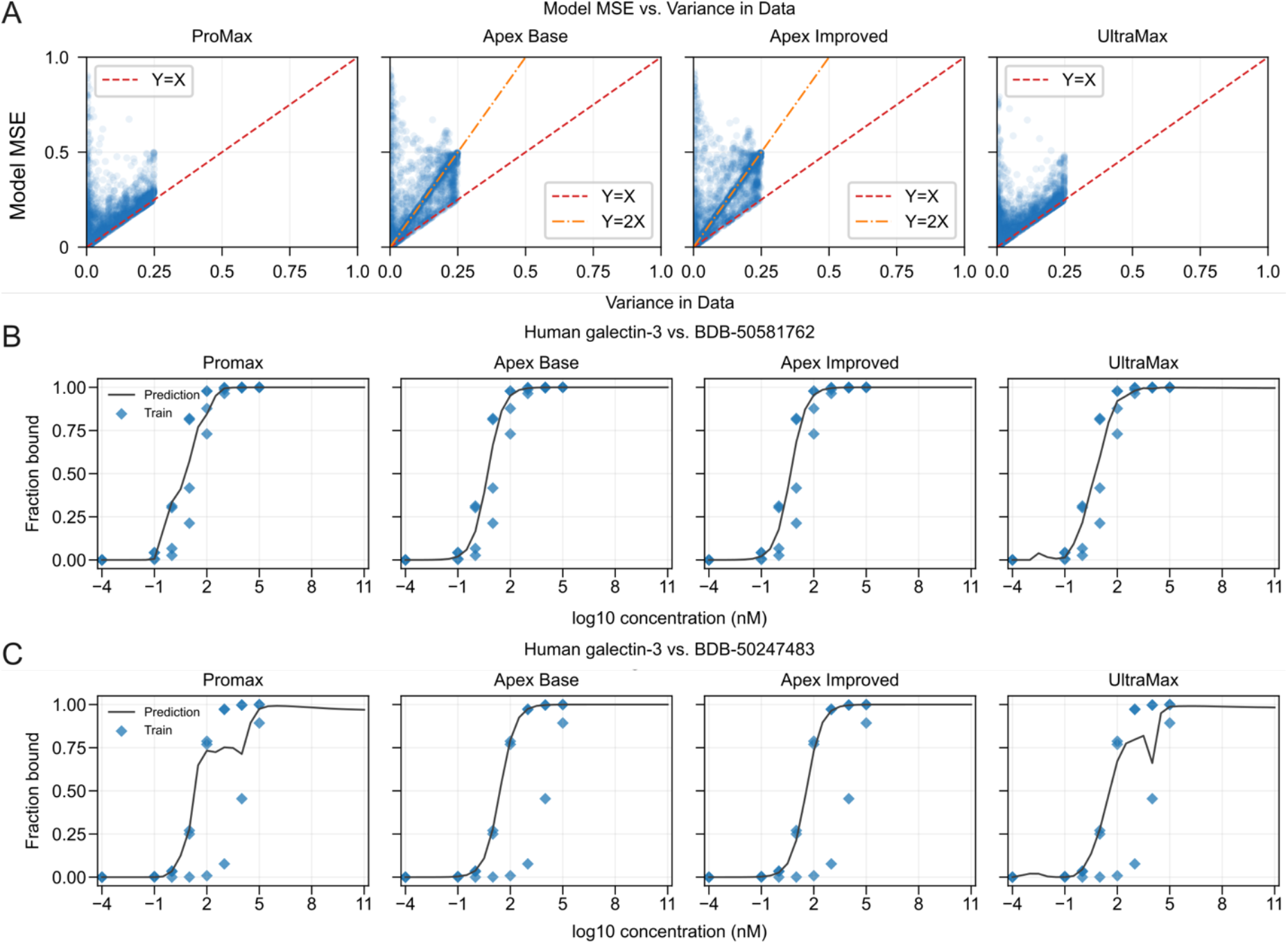
Per-configuration errors, and concentration response. (A) Variance of fraction bound values for a given molecule, protein, and concentration configuration (blue dots) vs. model MSE for the same configurations. Entries with only one value of fraction bound for a configuration are not plotted. For clarity, we only show dots where the variance in the data exceeds 0.0001 (*n* = 3,626 from the 68,492 unique (molecule, protein, concentration) configurations in the test set). For ProMax and UltraMax models, many of the dots lie close to the lower bound, suggesting that the models perform well on entries that have multiple fraction bound values. For APEX Base and Improved variants, a population of dots lie on the *y* = 2*x* line (orange line with alternating dashes and dots). (B, C) Fraction bound as a function of log-concentration (in nM) for different molecules against human galectin-3. Training data are semi-transparent points; when overlapped, they appear darker. The behaviour of the ProMax and the UltraMax models are not always monotonically increasing, as a function of concentration. The APEX Base and APEX Improved models have this behaviour explicitly encoded in them but perform worse in general (Fig. 3).

As shown in Fig. 5, the predicted fraction-bound profiles differ substantially across models, proteins, and datasets. On the BindingDB subsets (upper portions of panels A and B), all four models recover the overall response structure reasonably well, with the Galectin-1 plots showing a clear sigmoidal increase and the Galectin-9 plots remaining concentrated near low binding values except for a small strong-binding tail (which UltraMax overpredicts). In contrast, the CFG subsets (lower halves of panels A and B) are more challenging. In these panels, APEX produces predictions that remain close to zero for much of the molecule range, whereas ProMax and UltraMax more broadly assign stronger-binding responses. Notably, for Galectin-1, UltraMax provides the closest overall match to the observed distribution, capturing the high-binding region better than ProMax and avoiding the overly suppressed profile produced by APEX. Overall, the figure highlights a clear difference between performance on seen BindingDB data and the more difficult CFG setting, where hidden-sample generalization becomes the dominant challenge.

**Fig. 5.**
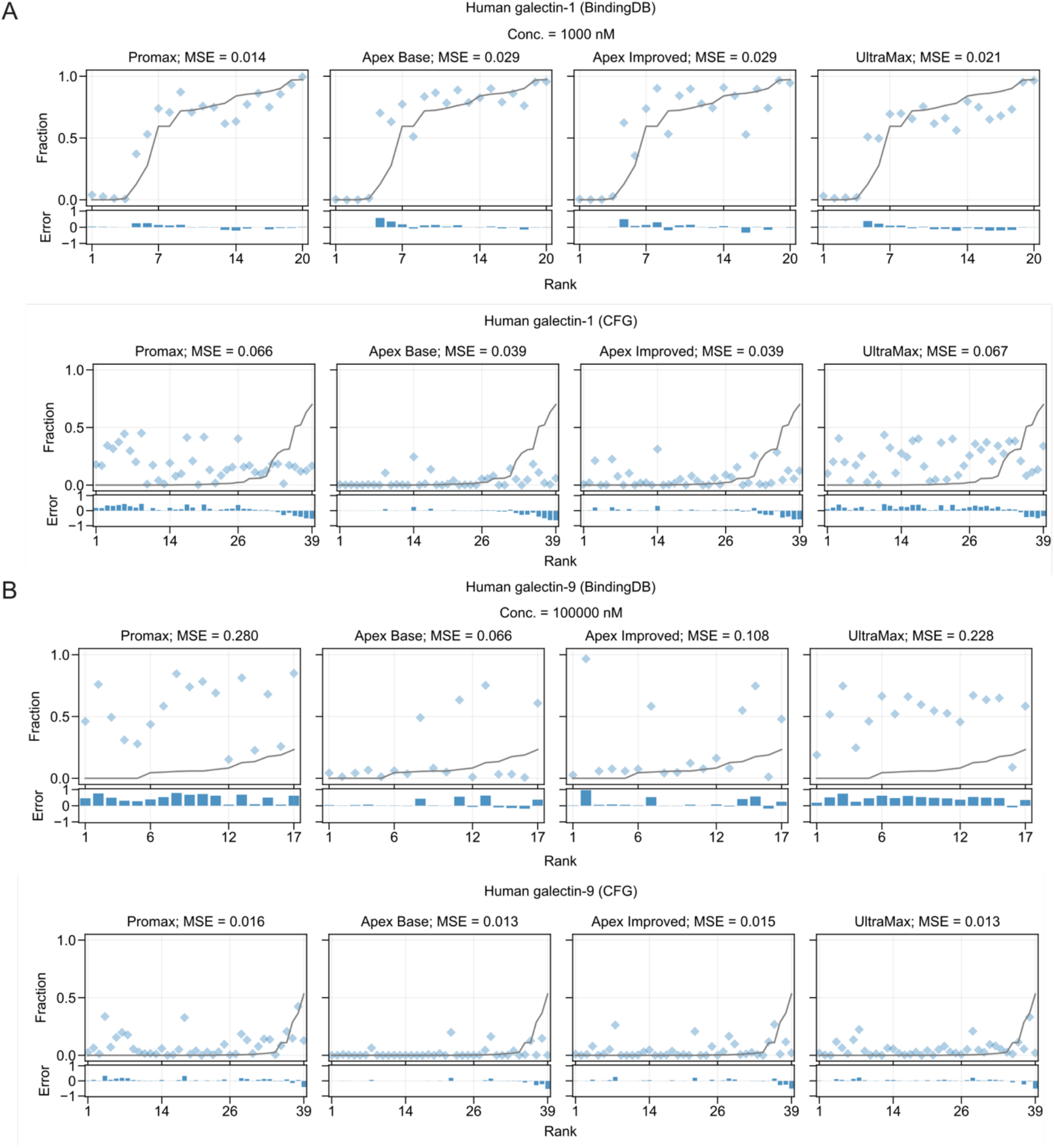
Fraction bound predictions of different models for proteins. (A) Human galectin-1 (BDB-0021/CFG-prot-1000013) and (B) Human galectin-9 (BDB-0025/CFG-prot-1002733) against different (molecule, concentration) configurations. The black line represents the data and diamonds represent predictions. For BindingDB proteins, we plotted subsets of the data that were at a particular concentration (1 and 100 µM respectively). For CFG proteins, we only plotted data at the maximum available concentration. The rank corresponds to the molecule rank when sorted in the increasing order of fraction bound values.

### Sample predictions on inversion of chirality

The MolFormer and MolCLR encodings are chiral-aware; they produce different encodings when different chiral centres of the same molecules are inverted. To test whether these models are truly chirality-aware, we tested them using a fundamental knowledge of chirality in protein-ligand interactions: starting from a strong binding chiral ligand L for protein P and inverting the chirality of all stereocenters creates a mirror image molecule ⅃. Strength of binding between ⅃and P is significantly reduced when compared to L-P interactions. Decreased interactions between a protein and a mirror-image of its native ligand can be traced as far back as 1858 publications of Louis Pasteur ^30,31^. These observations have really been solidified by 1990’s with mirror-image protein synthesis ^32^ and mirror-image phage display ^33^ and popularized in recent discussions of mirror-image life forms ^34,35^. To create “mirror image” ligands, we elect to use ligands that have five or more stereocenters to ensure that at least one of these stereocenters occurs at the critical interface between a protein and a ligand.

We augmented the test set by inverting all the chiral centers of molecules; the ground truth for these cases were set to zero binding, consistent with the expectation that proteins do not normally have substantial interactions with enantiomers of their regular substrates. We observe that these mirror-molecules are predicted to have different bindings than the original ones, however the distribution of these binding events for native and mirror ligands are very similar in many cases (**Figure 6**). The mirror-inversion test made it clear that models lack the necessary sensitivity to properly handle the stereochemistry in these augmented test cases. This confirms that the model bases its predictions on non-chiral cues and insufficiently utilizes the chiral aspects of molecules – key determinants of binding.

**Fig. 6.**
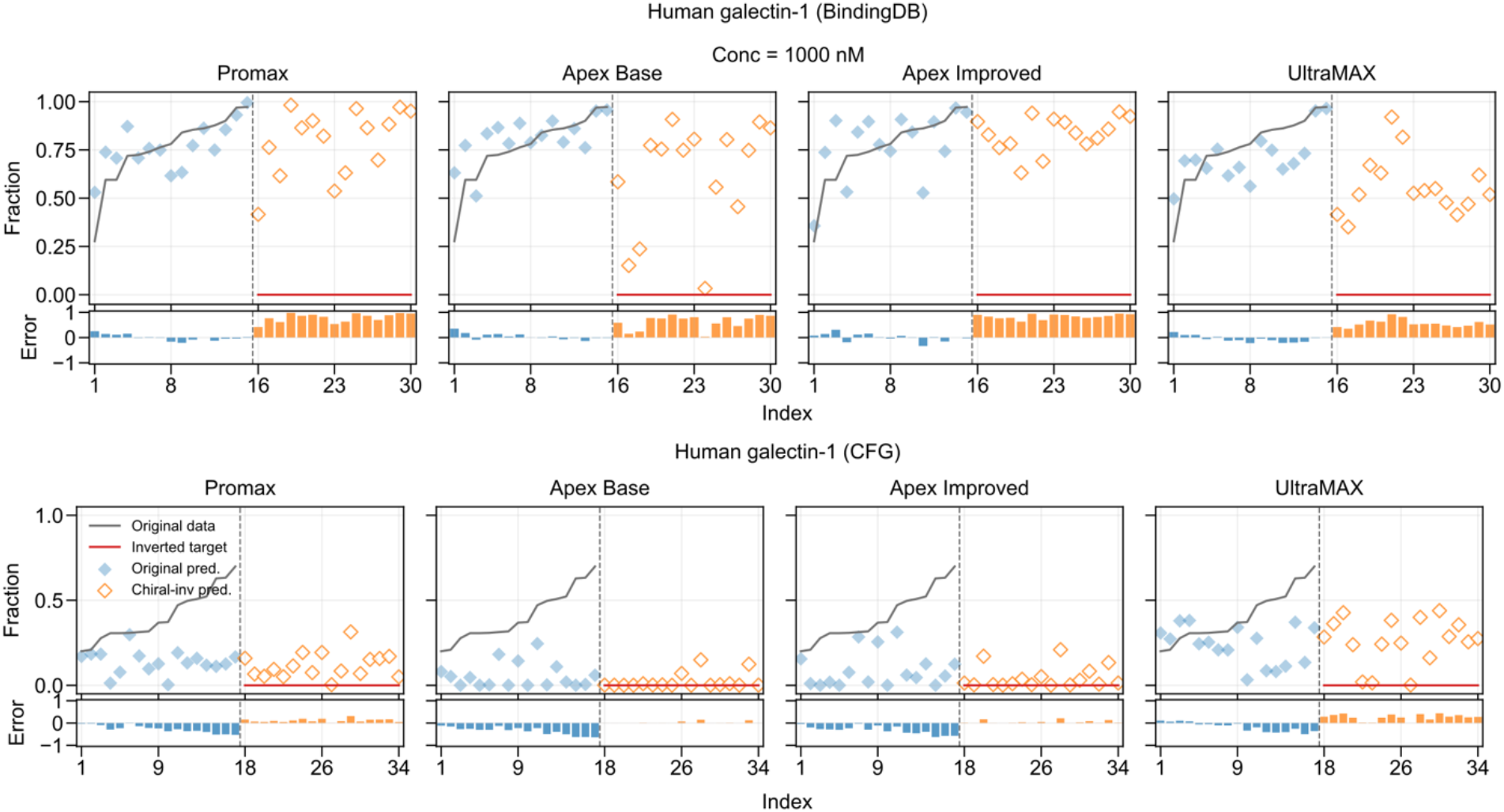
Fraction bound predictions on molecules of inverted chirality across the different models. We show predictions for human galectin-1 at a concentration of 1000 nM for the BindingDB examples and at the highest available concentration for CFG examples. The models predict similar values of fraction bound for the augmented molecules as the original test set (hollow orange diamonds).

## Discussion

In this work, we investigated glycan–protein interaction prediction as a multimodal regression problem using pretrained protein embeddings (ESM-2) with complementary glycan/molecule representations (MolFormer, MolCLR, ETKDGv3 derived). The use of loss functions tailored to bias against the tail of non-binding cases was critical for producing models that were able to capture rare strong-binding interactions. Rarity of interactions and class imbalance is a fundamental property of all protein-ligand interactions; however, only the CFG dataset offered a reasonable example of this imbalance. Almost every glycan on CFG arrays was tested in binding to almost every GBP. The non-binding data in the glycan microarray repository has not been discarded or censored as “useless”; it offered experimentally-validated negative data. In contrast, BindingDB is severely biased towards reporting of “good binders”. This bias has been discussed in medicinal chemistry literature for decades ^36,37^. The problem of *censorship of the observations that do not work* is now apparent in many fields of chemistry: for example, chemical databases exhibit a profound reporting bias, primarily emphasizing high-yielding reaction instances ^38,39^. There is a documented criticism of inflation of reaction yields ^40^ and we observe similar bias towards reporting of strong binders in publications that describe molecular discovery. Peer-reviewed literature has a bias against reporting weak binders and binders that have no “important biological activity”. To understand fundamental importance of compounds that simply bind targets rather than elicit *some* biological activity, we recommend a 2018 editorial from Stuart Schreiber ^41^. What may or may not be clear in Schreiber’s editorial is that “binding” is not a singular trait attributable to singular protein but rather a proteome-wide set of traits confirming that “interactions of a binder L and a protein P occurs only with protein P and it does not occur with other proteins in the same organism”. We refine Schreiber’s vision as: future must focus on the discovery of compounds that not only bind to a desired protein but also have proven lack of binding to many other proteins. By embracing the need for “non-binding data”, the severe imbalance of binding datasets and rarity of binding becomes a goal rather than inconvenience.

At present, we do not understand why the small-molecule cases train better than the glycan data. One hypothesis is that suboptimal learning is due the class imbalance between the two in the training set. In the future, to achieve a better balance between the two, future modellers face two options: Option 1: rejecting more of the BindingDB data, at the risk of the resulting model being impaired by a smaller chemical background, or option 2 the harder task of adding more glycan-binding data. A completely independent explanation is that difference arises from the higher numbers of chiral centres in glycan data and scarce examples of chirality-dependent interactions in small molecule datasets; in turn, chirality has been previously reported as a hard-to-learn feature ^11^. In a simple mirror-inversion of strong binders, we failed to see the expected loss of binding in the models. This observation strengthens the hypothesis that chirality, an important molecular feature of glycans, is not being learned to full effect by the models. A future direction may be to explore training approaches intended to strengthen the models’ responses to stereochemical aspects of the molecules, such as by augmenting the training set with enantiomers of strong binders, which are expected to be weak binders.

Our results show that multimodal representations of molecules can produce usable models, with UltraMax, our model incorporating the most modalities, achieving the best accuracy. We cautious that ETKDG-modelled inter-atomic “distances” ^27^ are unlikely to have any relation to the dynamic distances between atoms in a Boltzmann ensemble of molecular conformers. Still, such “distances” provide some important information that augments the learning process. Future paths should test the simplification of the models to explore whether the complexity is warranted through approaches like systematic ablation studies and better exploring the impacts of the different modalities used by the models and their fusion.

We compared prediction of fraction bound to prediction of *K*_D_ values and *K*_D_-surrogates calculated from microarray data. APEX models that use such “*K*_D_” values performed poorly compared to the other models. Equation-based models of binding interactions such as Hill, Langmuir adsorption, Hendersson-Hasselbach, etc. always have underlying assumptions and yield incorrect predictions if the stringent assumptions are violated. Although the use of equations has been used in modeling of protein-glycan interactions (8), the cooperative and multivalent aspects in glycan-protein binding appear significantly more frequently than in the interactions of drug-like small molecules to proteins. Calculation of *f* from reported *K*_D_ for small-molecule interactions is a generally safe whereas conversion of *f* from arbitrary binding observations to *K*_D_ is not. In development of ML models that aim to predict ligands protein interactions from real-world observations, a seductive nature of single *K*_D_-like value as catch-all descriptor must be carefully evaluated. We strongly believe that future ML models must account for the fundamental aspect of protein-glycan interactions across the range of glycan concentrations. We particularly urge ML practitioners to focus on near-saturation concentrations of glycans that occur in crowded biological environments like in glycocalyx and learn from century of observation of effect of near-saturated concentration of glycans in areas like vitrification, cryobiology, protein preservation and fundamental role of solvation in biology ^42-44^. ML models that learn from binding events at specific concentrations are likely to incorporate more fundamental knowledge when compared to models that predict “binder/not-a-binder” classes or single binding parameters.

The task in this manuscript always employs only one concentration and the use of a single concentration in the interaction that takes place between soluble ligand and soluble protein may be counterintuitive. The use of single concentration is key to data integration because only one concentration *exists* in experiments that measure binding of soluble protein P to immobilized ligands L such as glycans present on a glycan array or glycocalyx on the surface of living cells. A model of 1:1 monovalent binding equilibrium between soluble P and L can be solved explicitly to demonstrate that only one concentration is necessary and sufficient to compute the fraction bound ^10^. Analysis based on a 1:1 monovalent model is a limitation because many protein glycan interactions are multivalent and cooperative ^45,46^. We believe that it is possible to prove that in any protein-ligand interaction, whether monovalent or multivalent, there exists a universal local concentration of ligand in a coordinate system associated with protein which would be necessary and sufficient to calculate the fraction bound. Effective molarities employed in analysis of multi-valent ligands ^47^ or ligands tethered to the surface of proteins ^48^ are conceptually similar to the “local concentration”. This reduction from a multi-concentration framework to a single concentration framework is reminiscent of the reduction of any two-body problem to a one-body problem in classical mechanics ^49^. We are presently uncertain how data from multivalent and cooperative multi-step binding events between ligands and protein can be integrated into machine learning models that predict the strength of protein-ligand interactions; however, we are certain that data from the field of glycan-protein recognition will offer rich examples of such interactions.

## Code availability

The datasets and the code used to train the models are available for download at https://doi.org/10.5281/zenodo.20758183.

## Acknowledgments/Disclosure

This work originated as a course project for CMPUT 469 University of Alberta, portions of this work have previously been submitted for academic credit by some of the co-authors. We would like to acknowledge that this work used services and support provided by CFI-MSI GlycoNet Integrated Services, Canadian Institutes of Health Research #180445 (RD); Natural Sciences and Engineering Research Council (Canada) Discovery Grant #RGPIN-2016-402511 (R.D.) Genome Canada (R.D.); Canadian Foundation for Innovation (R.D.); Natural Sciences and Engineering Research Council (Canada) Discovery Grant #RGPIN-2019-04927 (R.G.), and CIFAR AI Chair (R.G.).

## Author contributions

Conceptualization: E.J.C., R.G., R.D.; Data Curation: E.J.C., S.H., Methodology: H.Y., W.L., W.Z., Z.C.; Formal analysis: H.Y., W.L., W.Z., Z.C., A.S., E.J.C., R.G., R.D.; Supervision: R.G., R.D.; Visualization: H.Y., W.L., W.Z., Z.C., A.S., E.J.C.; Writing - original draft: H.Y., W.L., W.Z., Z.C., A.S., E.J.C.

## Notes

https://doi.org/10.5281/zenodo.20758183

### Competing Interest Statement

The authors have declared no competing interest.

### Summary of Updates

removal of word breaking hyphen from online title and abstract

