## Supplemental Text and Figures for "Development of Deep-Learning Models that Predict Quantitative Protein-Ligand Interactions in Glycobiology as a part of a Capstone Course"

### Supplemental Material for Deep-Learning Resources for Studying Quantitative Glycan-Mediated Protein-Ligand Interactions

#### Methodology

Predicting glycan–protein interactions on our dataset requires handling a severe label imbalance. This label imbalance arises because many of the molecule-protein pairs in the dataset have low fraction-bounds (do not bind); this is especially true at lower protein concentrations.

To address these requirements, we develop an end-to-end deep learning pipeline consisting of three stages: data preprocessing, multimodal feature extraction, and interaction modeling with progressively more expressive architectures. Tail-aware objective functions are further introduced to handle the long-tailed distribution of binding data (see Fig. 1E).

Chirality enters the model through both symbolic atom descriptors and geometry. The MolFormer and MolCLR encoders can encode stereochemistry when stereochemical markers are present in the SMILES string. UltraMax makes such signals more explicit in the structural branches: during SMILES string parsing, RDKit assigns stereochemistry, and each atom feature vector includes a chiral-tag feature and a Cahn–Ingold–Prelog R/S-annotation where possible. The same atom-feature tensor is used by both the bond-graph branch and the distance-aware branch. In addition, the generated conformer can reflect stereochemistry geometrically, because different stereochemical assignments can lead to different spatial arrangements of atoms.

The models are implemented using the PyTorch library. Protein features are obtained from ESM-2 embeddings, while glycans are represented using MolFormer embeddings [1], MolCLR molecular topology-aware embeddings [2], and ETKDG-modelled inter-atomic distances [3]. Additional standard scientific libraries were used for data processing and evaluation; these are:

RDKit https://www.rdkit.org
ESM
Transformers Transformers: State-of-the-Art Natural Language Processing (Wolf et al., EMNLP 2020) https://aclanthology.org/2020.emnlp-demos.6/
https://github.com/huggingface/transformers
PyTorch https://pytorch.org
PyTorch Geometric https://pyg.org
PyTorch Scatter https://github.com/rusty1s/pytorch_scatter
scikit-learn https://scikit-learn.org
NetworkX https://networkx.org
Pandas https://pandas.pydata.org
NumPy https://numpy.org
SciPy https://scipy.org
Matplotlib https://matplotlib.org
tqdm https://tqdm.github.io

**ProMax: Transformer Based SMILES Encoding and Explicit Atom-Level Graph Encoding**

ProMax is our core multimodal baseline model for glycan–protein interaction prediction. This model integrates inputs of molecular structure, protein sequence and concentration through modality-specific encoders, followed by an explicit stage for interaction modeling and protein concentration-dependent prediction. Protein sequences are encoded using pretrained ESM-2 embeddings (esm2_t33_650M_UR50D), while molecules are represented through two views that are expected to have complementary emphases; MolFormer embeddings which are hoped may contain global molecular contexts and MolCLR embeddings which are derived from molecular bond-structure graphs with their explicit local information. These molecular representations are projected into a shared latent space and fused using a learned gated fusion mechanism.

Fig. 2A provides a high-level overview of the ProMax architecture. As shown in the figure, the model begins with three input branches corresponding to protein sequence, molecular structure, and protein concentration. The encoded features are then passed to two parallel interaction modules—a cross-attention pathway and a low-rank interaction pathway—which together capture complementary aspects of protein–molecule binding.

MolFormer reads SMILES strings and has been trained to recognize various structural patterns within these, producing latent graph embeddings which contain its only signals of molecular structure. This learned process is not guaranteed to be exact and features such as the nesting of branch parenthesis may not be interpreted correctly (Fig. S1A). Similarly, there is no mechanism, except the training process to ensure that even such apparently simple tasks as evaluating a structure in the opposite direction (e.g. the esters -C(=O)OC- vs. -COC(=O)-) produce the same results. Here MolFormer-XL-both-10pct was used, with the last hidden states mean-pooled to produce 768-dimensional embeddings. In contrast, tools such as MolCLR parse SMILES strings and explicitly build the exact atom and bond network for each molecule before processing those graphs directly. In these representations, each molecule is represented as an explicit graph of atoms (nodes) joined by bonds (edges), where each of the nodes is labelled with chemically meaningful features such as the element, hybridization state, number of bonds, charge, and whether it is in a ring. Similarly bond type can be used to label the edges. In this work the MolCLR GIN encoder was used to produce 256-dimensional embeddings of the molecules which are expected to capture local structural contexts. Compared with the MolFormer-style pathway, this design begins the encoding with an explicit representation of the molecular structure, which is expected to be more chemically grounded by operating directly on atoms and bonds and ensuring that the actual molecule is available to the system.

ProMax combines two interaction pathways. A cross-attention module allows the fused molecule representation to attend over residue-level protein features, capturing localized binding-relevant regions, while a low-rank interaction branch models multiplicative relationships between protein and molecule representations. In addition, concentration is encoded by a small multi-layer perceptron (MLP) and used to modulate the interaction representation through learned gating and FiLM-style transformations [4] which combine the embeddings, allowing the model to account for experimental conditions when predicting binding. Although these conditioning operations are abstracted for clarity in Fig. 2, they are part of the full prediction pipeline.

The resulting multimodal representation is passed to an MLP prediction head to directly regress the observed fraction bound.

#### APEX

Instead of directly regressing on *f*, the fraction bound, the APEX models predict a latent value $\log EC_{50}$, which is then transformed into the predicted fraction bound using the Henderson-Hasselbach equation

$$f = \frac{1}{1 + \left(^{EC_{50}}/_{c}\right)} = \left(1 + 10^{\log EC_{50}}/c\right)^{-1}$$

where *c* denotes the concentration (see also Fig. S2). This fixed transformation is consistent with a particular model of biochemical binding.

Predicting $EC_{50}$ provides a concentration-independent representation of affinity, potentially reducing overfitting to dataset-specific conditions. This constraint on the model's behaviour, forces a behaviour that matches the expected one, and may prevent the model from learning spurious correlations while improving robustness on unseen molecule–protein pairs.

Because *f* is derived from $EC_{50}$ through a nonlinear mapping, prediction errors propagate unevenly across the binding curve. Small deviations in $\log EC_{50}$ can lead to larger changes in *f* in mid-affinity regions, while errors are naturally attenuated in the saturation regimes where $f \approx 0$ or $f \approx 1$. As a result, APEX achieves more stable and physically consistent predictions in low- and high-binding regimes, where errors are naturally compressed.

Two versions of APEX models were developed and results for both are presented. The original version was named APEX Base and the revised version APEX Improved. This later version uses protein-molecule attention heads to fuse the inputs. In training these heads a contrastive learning approach was adopted.

#### UltraMax: Multi-View Structural Fusion

UltraMax extends the earlier Promax and APEX models. It keeps the same ESM-2 protein representation and the transformer-parsed SMILES string representation from MolFormer, while strengthening the structural and interaction parts of the glycan-protein model. Its main additions are: (i) it's own explicit atom-level molecular structure encoder instead of the MolCLR signal, (ii) a complementary inter-atom distance-aware branch based on an RDKit predicted conformer, (iii) a learned multi-view gate that reasons jointly over the MolFormer, the bond-structure, and the distance based glycan evidence, and (iv) a deeper bidirectional cross-attention module for protein-glycan fusion. Fig. 2B summarizes this overall architecture.

The SMILES string branch processes the SMILES strings using a precomputed model (MolFormer) and uses its output as a molecular embedding. The molecular structure branch operates on the molecular connectivity graph (see Fig. S3A), where atoms are nodes and bonds define edges, so it captures local features of topology, branching, ring membership, and bond-local context, but does not have explicit coordinates. The distance-aware branch uses the same atom-level graph together with spatial distances from a predicted conformer, so it should better capture distance-dependent spatial relationships between atoms (Fig. S3C).

**Explicit Atom-level Molecular Structure Encoding**

A key departure from the earlier models is that UltraMax no longer relies on the mean-pooled atom neighbourhood embeddings produced by MolCLR's graph processing as a molecular structural signal. Instead, it uses RDKit to parse the SMILES string into an atom-level graph for each molecule and processes those graphs through its own pipeline. Each atom is labelled with several chemically important features. They are atomic number, attached hydrogen count, hybridization state, chiral tag, Cahn–Ingold–Prelog R/S labels, aromatic flags, ring membership, number of bound neighbours, and formal charge. These nine features can be mathematically represented as a matrix of $Feature_{molecule} \in \mathbb{R}^{N\times 9}$ , where N is the number of atoms in a molecule.

The structural connectivity of this graph is given in an adjacency matrix, $A_{molecule} \in \mathbb{R}^{N\times N}$ (Fig. S3B). This matrix $A_{molecule}$ serves as the foundation for the 2D encoder, as it represents the relationship between any two atoms $i$ and $j$ is represented by a binary value:

$A_{i,j} = 1$ a direct chemical bond exists between the two atoms, or

$A_{i,j} = 0$ no direct bond exists.

The bond-network encoder applies adjacency-constrained graph-attention layers over this matrix, allowing nodes in the model to aggregate information only from immediate neighbors. We use the convention that the diagonal entries $A_{i,i}$ are set to 1, creating self-loops, which allows atoms to retain their own features during the message-passing process. Together with the atom-level features, the bond-network representation can be expressed as $G = (Feature_{molecule}, A_{molecule})$

Because message passing follows the bond-graph, this pathway mainly captures local topological context. This includes bonding patterns, branching structure, ring membership, and stereochemically annotated atomic environments. While this branch is highly effective at representing how the glycan is connected, the binary nature of the adjacency matrix means it does not distinguish between different bond types (lengths) or capture spatial proximity between atoms that are not directly bonded; however those relationships may instead be captured by the distance-aware branch (Fig. S3D).

**Distance-aware 3D glycan encoding**

UltraMax integrates a 3D structural branch that utilizes the same atomic level features, $Feature_{molecule}$; but incorporates Cartesian coordinates ($x$, $y$, $z$). Concretely, for each molecule, RDKit's ETKDGv3 model [3, 5] is used to generate a plausible conformer with coordinates, $C \in R^{B\times N\max\times 3}$, for each atom in physical space, and the physical-distance branch associates close-pairings (under 6.0 Å) in these coordinates with an atom feature matrix and performs distance-aware message passing. Unlike the bond-based molecular graphs, which exchanges information only between chemically bound atoms, this distance-aware branch employs a 6.0 Å radius cutoff. Atoms are neighbours, if their centres are within 6 Å of each other, which allows atoms that are distant in the molecular-structure, but close in physical space to exchange information through geometric message passing. To represent these spatial relationships, the model calculates a pairwise 3D distance matrix (Fig. S4D) where each Euclidean distance is encoded using eight channel Radial Basis Function (RBF) networks[6]. This distance-aware approach enables UltraMax to capture aspects of steric proximity and conformational organization—spatial cues that purely topological encoders cannot directly capture—providing a physically grounded description of the glycan-protein interaction. Internally, this branch uses a hidden width of 256 nodes, 8 attention heads, and a distance encoding with 8 channels. Because this physical-distance encoder aggregates across atoms that are spatially close even when they are not directly adjacent in the molecule, UltraMax should capture not only the molecular topology, but also aspects of molecule's spatial geometry, enabling the model to represent steric and conformational cues that are inaccessible to purely sequence-based or to structural encoders.

A significant limitation of this approach is that to perfectly represent inter-atomic distances within the flexible shapes of the glycans and other molecules requires ensembles of conformers to completely sample the state-space. While this is approximated with only a single example conformer, it is obviously an incomplete representation, but it still contains information about the relative positions of atoms, especially those nearest to each other, under the 6 Å cutoff. We expect that even this single conformer may serve as a hint to the model, perhaps helping to guide it to better predictions, but with the caveat that a single conformer is not a full description of the spatial arrangement of the atoms. We note that extending this to a full treatment of glycan/molecule conformational space is a non-trivial problem, e.g. Refs.[7-9] and that in other work, conformers have been reported to be informative[10].

**Gated Multi-view Fusion**

Beyond adding the physical-distance branch, UltraMax also generalizes the gated learning mechanism by learning jointly over the MolCLR embedding, the explicit bond-based molecular topology, and the physical-distance molecular encodings (Fig. 3B). Instead of selecting between two representations, this model computes the three separately and then applies a learned gate to assign sample-specific soft weights across these three modalities. To stabilize training, the gate is initialized to begin from an approximately uniform distribution and gradually transitions toward a learned modality preference, allowing the model to adaptively determine to what extent each of the three molecular views is informative for each given interaction. Let $H_1, H_2, H_3$ denote the projected branch summary. The model concatenates the three branch summaries, represented as $z = [H_1;\ H_2;\ H_3]$. The weight for each branch is learned as in the following equation: $W = softmax(MLP(z))$, where $W \in \mathbb{R}^3$. Each $H_i$ branch is then scaled by their learned weight $w_i \in W$.

To further compare the difference between the bond-based and distance-based representations, and learn a precise weight to fuse them, we formulated a comparison representation D, using the raw states, their element-wise product, and their absolute difference.

$$D = [H_{\text{bond}}, H_{\text{distance}}, H_{\text{bond}} \odot H_{\text{distance}}, |H_{bond} - H_{\text{distancce}}|\,]$$

The purpose of *D* is to give the fusion module a direct comparison between the bond- and distance-based representations at each atom. When the two are consistent, the product term highlights their agreement; when they differ, the absolute-difference term exposes the mismatch, helping the model decide whether to rely more on the bond-network, inter-atom distances, or a learned combination of both.

Finally, A sigmoid gate decides how much each atom-feature dimension should lean toward the bond-network branch versus the conformer distance-network branch.

$$G = \sigma(Linear(LayerNorm(D)))$$

The total fused representation is shown by the following equation:

$$H_{fuse} = G \odot H_{\text{bond}} + (1 - G) \odot H_{\text{distance}} + H_{\text{SMILES}}$$

#### Cross-attention and prediction

After the molecular-topology and physical-distance embeddings are fused they are combined with the global MolCLR embedding to form the final molecular representation used downstream for interaction modeling: $H_{molecule} = [\, H_{\text{MolCLR}} \,;\, H_{fuse} \,]$. In this case, $H_{\text{MolCLR}}$ is expected to serve as a global summary of the molecule, and $H_{fuse}$ provided a refined description of it. Before passing into cross attention, $H_{molecule}$ was first passed through a self-attention layer. In parallel, the other major input, the protein representation, $H_{protein}$ was also passed through its self-attention layer and projected into a shared space. Then both protein and molecules were passed through 4 layers of a bidirectional block, each bidirectional block first updated the protein tokens using the glycan context, then updated the glycan tokens using the newly updated protein context. The process is expressed using the following two equations.

$$\widetilde{H}_{protein} = CrossAttn\left(H_{protein}, H_{molecule}\right)$$

$$\widetilde{H}_{molecule} = CrossAttn\left(\widetilde{H}_{protein}, H_{molecule}\right)$$

Finally, the prediction is computed using MLP and sigmoid: $pred = sigmoid\left(MLP\left(\widetilde{H}_{protein}, \widetilde{H}_{molecule}, C\right)\right)$, where *C* is the concentration

#### Training Objective

The model was trained with AdamW using batch size 128, weight decay of $10^{-3}$, and gradient clipping at norm 1.0. The training was for up to 80 epochs with early stopping with a patience of 15 epochs. All linear weights were Xavier-initialized while biases were set to zero. The final output bias was set to the logit of the mean fraction bound in order to stabilize early predictions.

Because the response distribution was imbalanced weighted sampling was used. Fraction-bound values were divided across 10 bins, and each bin was assigned a sampling weight based on a mixture of the empirical label distribution and a uniform bin distribution. In the initial epochs, stronger weights sere used for underepresented regions, reducing with a cosine-schedule to closer to the actual training distribution. In addition, label distribution smoothing was used to assign continuous sample weights from the smoothed empirical target distribution.

**Objective Functions and Loss Designs: Long-tail Aware Processing**

Examination of the binding data shows that many of the molecule-protein pairs have low fraction-bounds (i.e. do not bind) especially at lower protein concentrations. To address this long-tail distribution of binding data, we adopt a unified loss formulation that combines several terms and is intended to emphasize high-affinity samples while maintaining stable optimization:

$$L_{total} = L_{main} + L_{soft\_tail} + L_{rank} \text{ (+ } L_{logEC} \text{ for APEX)}$$

Here the overall loss, $L_{total}$, is the sum of $L_{main}$ a conventional mean-squared error (MSE), $L_{soft\_tail}$ a sigmoidal weighting that emphasizes strongly-binding cases, and $L_{rank}$ which enforces correct ordering between predictions. For APEX, an additional $L_{log\ EC}$ term operates in the log $EC_{50}$ space to supervise the predictions.

The soft-tail weighting follows a logistic-function that increases smoothly with the ground-truth binding value,

$$L_{soft\_tail} = 2\ \sigma((\text{predicted value} - 0.4) / 0.08),$$

which biases the model to focus on the tail of high-affinity interactions without introducing discontinuities in the overall loss function. The $L_{rank}$ term is based on comparing the relative magnitudes of pairs of datapoints in both the ground truth and the predictions. It is given by

$$L_{rank} = \tfrac{1}{2}\, p_{ij} / N,$$

where $p_{ij}$ is the number of sampled pairs with the same order as in the ground truth, and N is the total number of pairs this test is evaluated over.

Unlike fixed reweighting schemes, all loss weights are adaptively adjusted based on the batch distribution, enabling dynamic emphasis on informative regions of the data.

This design tries to address a key limitation of prior approaches such as GlyNet, which rely on global error minimization and tend to underfit rare, but biologically important, strong binders. By combining tail-aware weighting with ranking supervision, our objective encourages both accurate prediction and correct ordering of high-affinity interactions, leading to improved performance in the most critical regions of the task. To ensure stable optimization under strong reweighting, we apply mask-aware attention to prevent padded tokens from contributing to gradients.

**Analysis of Loss Functions**

For a minibatch of size *N*, let $y_i$ denote the ground-truth fraction-bound value and $\hat{y}_i$ the prediction, where larger values are associated with stronger binding. Weighting values, $\lambda$, were set to 1.0, 0.8, and 0.3 respectively.

$$\mathcal{L}_{total} = \lambda_{MSE}\mathcal{L}_{MSE} + \lambda_{soft-tail}\mathcal{L}_{soft-tail} + \lambda_{rank}\mathcal{L}_{rank}$$

The soft-tail loss function is a weighted MSE; where for each pair, if the true binding value $y_i$ is high, then the $y_i - center$ will be high. It will assign more weight to the loss, to force the prediction to be accurate. center and scale of the logistic function are set to 0.4 and 0.08 respectively.

$$\mathcal{L}_{soft-tail} = \frac{1}{N}\sum_{i=1}^{N} w_i \cdot (\hat{y}_i - y_i)^2 \qquad w_i = \frac{logistic(y_i - center)}{scale}$$

$\mathcal{L}_{rank}$ is a loss term to enforce relative ordering of pairs of datapoints. For a minibatch size of *N,* i.e *N* prediction in total, we will have $C(N, 2)$ prediction pairs. We then select a subset of these pairs, P, where the difference between their true fraction bond is greater than a threshold $\delta$. In our case, $\delta = 0.1$.

$$P = \{ (i,j) : y_i - y_j > \delta \}$$

We then computed the pair wise loss: $\ell_{pair(i,j)} = max\left(0, m - \left(\hat{y}_i - \hat{y}_j\right)\right)$. In our case, *m*, the margin is 0.2.

$$\mathcal{L}_{rank} = \left(\frac{1}{|P|}\right)\sum\nolimits_{(i,j)\in P} \ell\_pair(i,j) \quad if\ |P| > 0,\ else\ 0$$

For the loss function in APEX models, an extra term $\mathcal{L}_{KD}$ incorporating a Huber loss was added. $\lambda_{KD}$ is 0.2

$$\mathcal{L}_{total} = \lambda_{MSE}\mathcal{L}_{MSE} + \lambda_{soft-tail}\mathcal{L}_{soft-tail} + \lambda_{rank}\mathcal{L}_{rank} + \lambda_{KD}\mathcal{L}_{KD}$$

$$\mathcal{L}_{KD} = \frac{1}{N}\sum_{i=1}^{N} Huber(\widetilde{KD}_i - KD_i)$$

$$Huber(\widetilde{KD}_i - KD_i)\begin{cases} 0.5 \times (\widetilde{KD}_i - KD_i)^2, & |\widetilde{KD}_i - KD_i| < 0.15 \\ 0.5 \times (|\widetilde{KD}_i - KD_i| - 0.75), & |\widetilde{KD}_i - KD_i| \geq 0.15 \end{cases}$$

#### Supplemental Figures

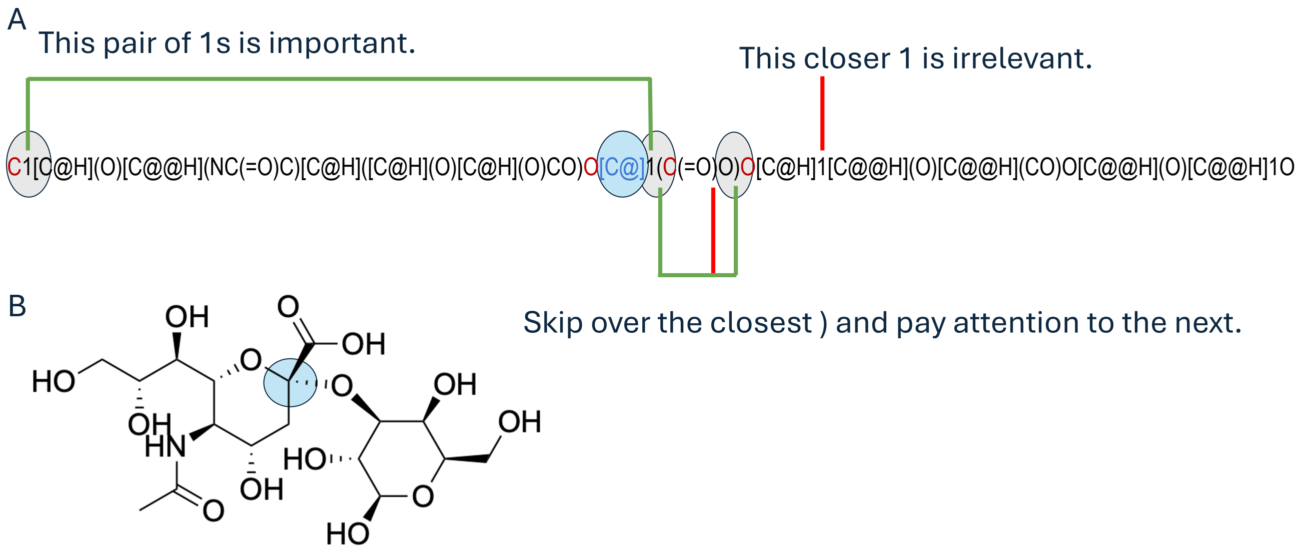


**Fig. S1** Complexity of interpreting SMILES strings. A) Determining neighbouring atoms (in red) of the atom in the blue circle is relatively complex and requires understanding the relationships between the different "1" characters in the string as well as the nested parenthesis pairs. B) In contrast a graph of the atoms and their bonds groups all the neighbours around the atom (blue circle) and the four neighbours are easily found.

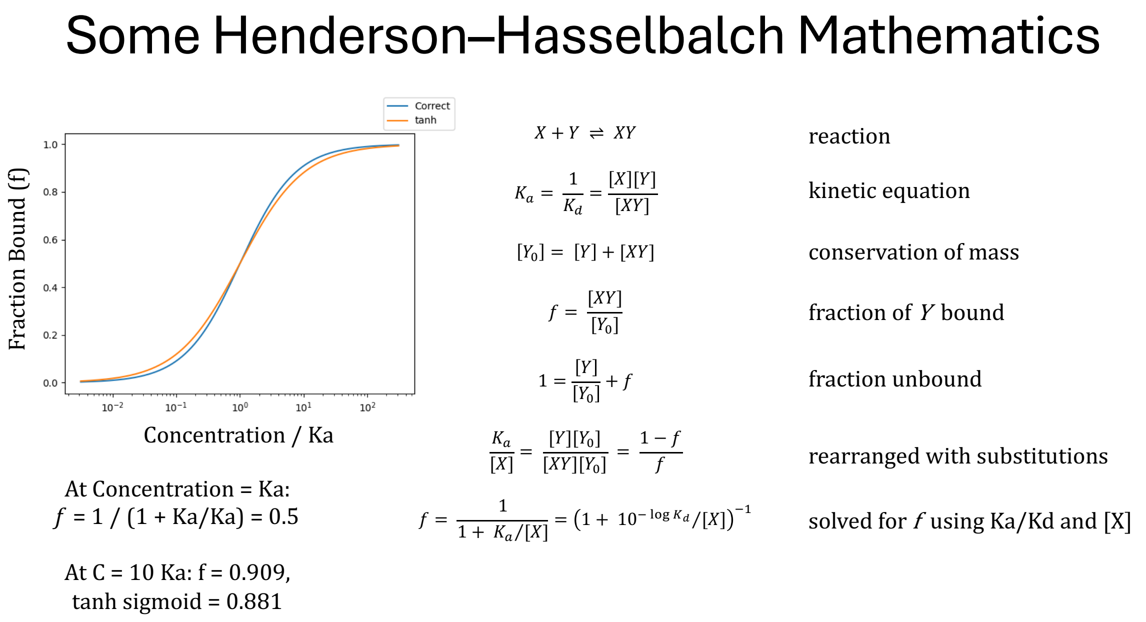


**Fig. S2** Mathematics of the Henderson-Hasselbalch equation.

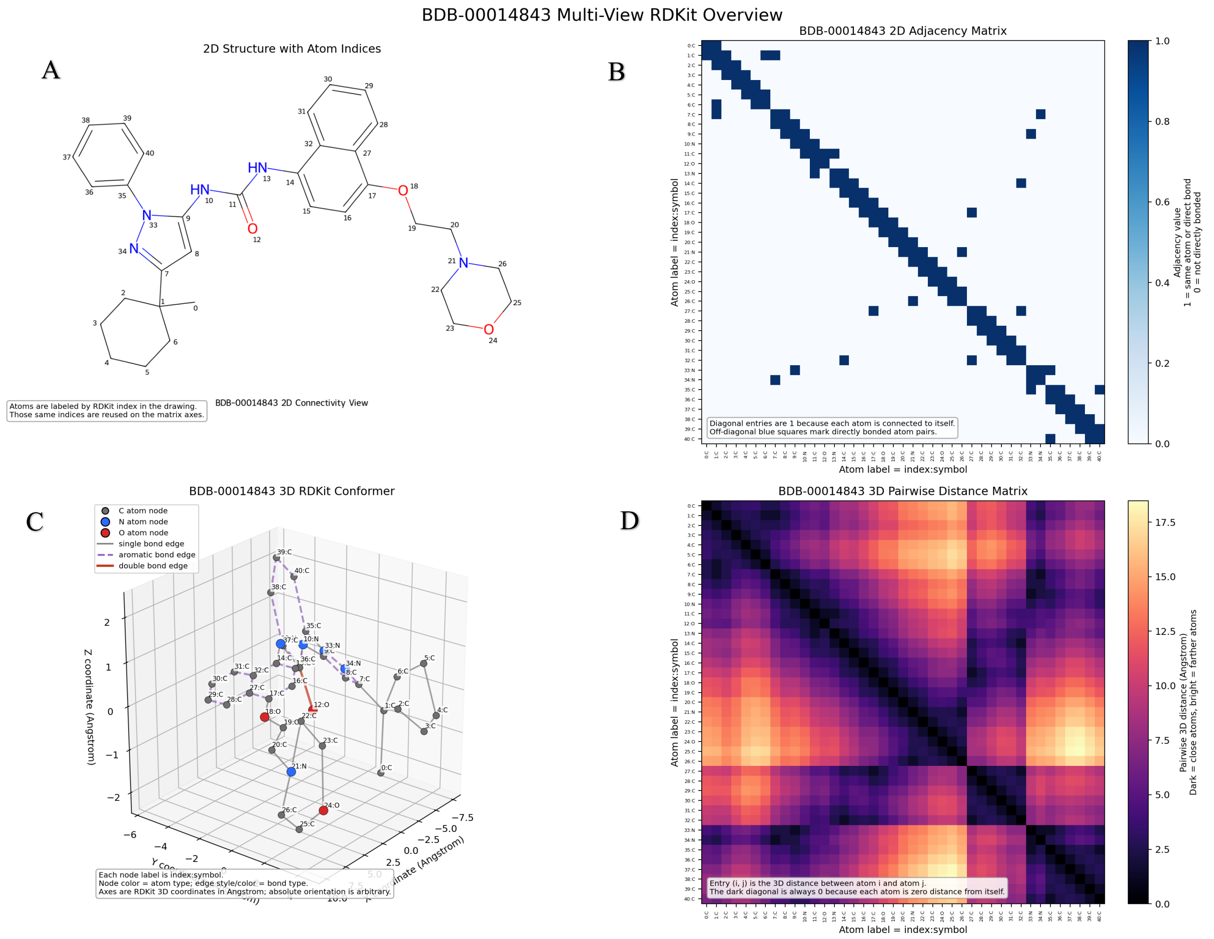


**Fig. S3** RDKit-based structural views for an example molecule (BDB-00014843). This figure illustrates how the bond-based and inter-atom distance-based structural branches represent the molecule. (A) The 2D chemical drawing with atom indices, which serve as reference labels for the matrix-based representations. (B) The 2D adjacency matrix utilized by the graph-attention encoder; diagonal entries are 1 (self-connectivity), and blue off-diagonal cells indicate directly bonded atom pairs. (C) The RDKit conformer used by the distance-aware branch, where node colors indicate atom types and Cartesian coordinates are measured in angstroms. (D) The pairwise distance matrix, where entry (i, j) represents the Euclidean distance between those atoms; darker values indicate spatial proximity while brighter values indicate greater distance.


1. J. Ross, B. Belgodere, V. Chenthamarakshan, I. Padhi, Y. Mroueh and P. Das, Nat. Mach. Intell., 2022, 4.
2. Y. Y. Wang, J. R. Wang, Z. L. Cao and A. B. Farimani, Nat. Mach. Intell., 2022, 4, 279–287.
3. S. Riniker and G. A. Landrum, J. Chem. Inf. Model., 2015, 55, 2562–2574.
4. E. Perez, F. Strub, H. d. Vries, V. Dumoulin and A. Courville, arxiv.org, 2017.
5. S. Wang, J. Witek, G. A. Landrum and S. Riniker, J. Chem. Inf. Model., 2020, 60, 2044–2058.
6. D. S. Broomhead and D. Lowe, Complex Systems, 1988, 2, 321–355.
7. S. H. Wang, T. J. Wu, C. W. Lee and J. Yu, J. Biomed. Sci., 2020, 27, 93.
8. T. J. Seifert, D. Kumar, M. Etzkorn, S. Rauschenbach, K. Kern, K. Anggara and U. Schlickum, arXiv.org, 2026.
9. I. L. Grothaus, G. Bussi and L. Colombi Ciacchi, J. Chem. Inf. Model., 2022, 62, 4992–5008.
10. L. Thomes, R. Joeres, Z. Akdeniz and D. Bojar, Nat Commun, 2025, 16, 11136.
